# A Distributed Network for Multimodal Experiential Representation of Concepts

**DOI:** 10.1101/2021.07.05.451188

**Authors:** Jia-Qing Tong, Jeffrey R. Binder, Colin J. Humphries, Stephen Mazurchuk, Lisa L. Conant, Leonardo Fernandino

**Author notes:** **Corresponding Author:** Leonardo Fernandino, Ph.D., Department of Neurology, Medical College of Wisconsin, 8701 W Watertown Plank Rd, Milwaukee, WI 53226.

## Abstract

Neuroimaging, neuropsychological, and psychophysical evidence indicates that concept retrieval selectively engages specific sensory and motor brain systems involved in the acquisition of the retrieved concept. However, it remains unclear which supramodal cortical regions contribute to this process and what kind of information they represent. Here, we used representational similarity analysis of two large fMRI data sets, with a searchlight approach, to generate a detailed map of human brain regions where the semantic similarity structure across individual lexical concepts can be reliably detected. We hypothesized that heteromodal cortical areas typically associated with the “default mode network” encode multimodal experiential information about concepts, consistent with their proposed role as cortical integration hubs. In two studies involving different sets of concepts and different participants (both sexes), we found a distributed, bihemispheric network engaged in concept representation, composed of high-level association areas in the anterior, lateral, and ventral temporal lobe; inferior parietal lobule; posterior cingulate gyrus and precuneus; and medial, dorsal, ventrolateral, and orbital prefrontal cortex. In both studies, a multimodal model combining sensory, motor, affective, and other types of experiential information explained significant variance in the neural similarity structure observed in these regions that was not explained by unimodal experiential models or by distributional semantics (i.e., word2vec similarity). These results indicate that, during concept retrieval, lexical concepts are represented across a vast expanse of high-level cortical regions, especially in the areas that make up the default mode network, and that these regions encode multimodal experiential information.

**Significance Statement:** Conceptual knowledge includes information acquired through various modalities of experience, such as visual, auditory, tactile, and emotional information. We investigated which brain regions encode mental representations that combine information from multiple modalities when participants think about the meaning of a word. We found that such representations are encoded across a widely distributed network of cortical areas in both hemispheres, including temporal, parietal, limbic, and prefrontal association areas. Several areas not traditionally associated with semantic cognition were also implicated. Our results indicate that the retrieval of conceptual knowledge during word comprehension relies on a much larger portion of the cerebral cortex than previously thought, and that multimodal experiential information is represented throughout the entire network.

## Introduction

Conceptual knowledge is essential for everyday thinking, planning, and communication, yet few details are known about where and how it is implemented in the brain. Assessments of patients with brain lesions have shown that deficits in the retrieval and use of conceptual knowledge can result from damage to parietal, temporal, or frontal cortical areas (Warrington and Shallice, 1984; Gainotti, 2000; Neininger and Pulvermüller, 2003; Damasio et al., 2004). Such neuropsychological findings have been extended by functional neuroimaging studies, which implicate a set of heteromodal cortical regions including the angular gyrus, the anterior, lateral, and ventral aspects of the temporal lobe, the inferior and superior frontal gyri, and the precuneus/posterior cingulate gyrus, all showing stronger activations in the left hemisphere (Binder et al., 2009; Hodgson et al., 2021). However, there is still debate on whether some of these areas – particularly the angular gyrus, the superior frontal gyrus, and the precuneus/posterior cingulate – are indeed involved in processing conceptual representations. It has been suggested, for example, that confounding factors such as task difficulty may be responsible for the angular gyrus activations found in the aforementioned studies (Humphreys et al., 2021).

Functional MRI (fMRI) studies also show that, in addition to these heteromodal areas, cortical areas involved in perceptual and motor processing are selectively activated when concepts related to the corresponding sensory-motor aspects of experience are retrieved (Meteyard and Vigliocco, 2008; Binder and Desai, 2011; Kiefer and Pulvermüller, 2012; Kemmerer, 2014). These findings are in agreement with “grounded” theories of concept representation, which predict that sensory-motor and affective representations involved in concept formation are re-activated during concept retrieval (Damasio, 1989; Barsalou, 2008; Glenberg et al., 2009).

Various models propose a central role for multimodal or supramodal hubs in concept processing, though both the anatomical location and information content encoded in these hubs remain unclear (Mahon and Caramazza, 2008; Binder and Desai, 2011; Lambon Ralph et al., 2017). One prominent theory postulates widespread and hierarchically organized convergence zones in multiple brain locations (Damasio, 1989; Mesulam, 1998; Meyer and Damasio, 2009). We previously proposed (Fernandino et al., 2016a) that these convergence zones are neurally implemented in the multimodal connectivity hubs identified with the “default mode network” (DMN) (Buckner et al., 2009; Sepulcre et al., 2012; Margulies et al., 2016). These areas closely correspond to those identified in a neuroimaging meta-analysis of semantic word processing (Binder et al., 2009). This idea is further supported by neuroimaging findings suggesting that the precuneus, posterior cingulate gyrus, angular gyrus, dorsomedial and ventrolateral prefrontal cortex, and lateral temporal areas encode multimodal information about the sensory-motor content of concepts (Bonner et al., 2013; Fernandino et al., 2016a, 2016b, 2022; Murphy et al., 2018).

In the present study, we used representational similarity analysis (RSA) with a whole-brain searchlight approach to identify cortical regions involved in multimodal conceptual representation. For a given set of stimuli (e.g., words), RSA measures the level of correspondence between the similarity structure (i.e., the set of all possible pairwise similarity distances) observed in the stimulus-related multivoxel activation patterns and the similarity structure for the same stimuli computed from an *a priori* representational model (Kriegeskorte et al., 2008). In contrast to previous RSA studies of concept representation, which used models based on taxonomic relations or word-co-occurrence statistics (Bruffaerts et al., 2013; Devereux et al., 2013; Anderson et al., 2015; Liuzzi et al., 2015; Martin et al., 2018; Carota et al., 2021), we used an experiential model of the information content of lexical semantic representations (henceforth, “conceptual content”) based on 65 sensory, motor, affective, and other experiential dimensions (Binder et al., 2016). Although not an exhaustive account of conceptual content, the experiential model addresses many relevant aspects of the phenomenological experience and the information content associated with lexical concepts. We used a searchlight approach (Kriegeskorte et al., 2006; Kriegeskorte and Bandettini, 2007) to generate a map of cortical regions where this multimodal experiential model predicted the neural similarity structure of hundreds of lexical concepts. Identical analyses were conducted on two large datasets to assess replication across different word sets and participant samples.

## Materials and Methods

### Participants

Participants in Experiment 1 were 39 right-handed, native English speakers (21 women, 18 men; mean age = 28.7; range: 20-41). Experiment 1 included data from 36 participants used in Fernandino et al. (2022), with 3 new participants added. Participants in Experiment 2 were 25 native English speakers 20 women, 5 men; mean age = 26; range: 19-40). None of the participants in Experiment 2 took part in Experiment 1. All participants in Experiments 1 and 2 were right-handed according to the Edinburgh handedness inventory (Oldfield, 1971) and had no history of neurological disease. Participants were compensated for their time and gave informed consent in conformity with a protocol approved by the Institutional Review Board of the Medical College of Wisconsin.

### Stimuli

The stimulus set used in Experiment 1 is described in detail in Fernandino et al. (2022). It consisted of 160 object nouns (40 each of animals, foods, tools, and vehicles) and 160 event nouns (40 each of social events, verbal events, non-verbal sound events, and negative events; mean length = 7.0 letters, mean log HAL frequency = 7.61). Stimuli in Experiment 2 consisted of 300 nouns (50 each of animals, body parts, food/plants, human traits, quantities, and tools; mean length = 6.3 letters, mean log HAL frequency = 7.83), 98 of which were also used in Experiment 1.

### Experiential Concept Features

Experiential representations for these words were available from a previous study in which ratings on 65 experiential domains were used to represent word meanings in a high-dimensional space (Binder et al., 2016). In brief, the experiential domains were selected based on known neural processing systems – such as color, shape, visual motion, touch, audition, motor control, and olfaction – as well as other fundamental aspects of experience whose neural substrates are less clearly understood, such as space, time, affect, reward, numerosity, and others. Ratings were collected using the crowd sourcing tool Amazon Mechanical Turk, in which volunteers rated the relevance of each experiential domain to a given concept on a 0-6 Likert scale. The value of each feature was represented by averaging ratings across participants. This feature set was highly effective at clustering concepts into canonical taxonomic categories (e.g., animals, plants, vehicles, occupations, etc.; Binder et al., 2016) and has been used successfully to decode fMRI activation patterns during sentence reading (Anderson et al., 2017, 2019).

### Procedures

In both experiments, words were presented visually in a fast event-related procedure with variable inter-stimulus intervals. The entire list was presented to each participant six times in a different pseudorandom order across 3 separate imaging sessions (2 presentations per session) on separate days.

On each trial, a noun was displayed in white font on a black background for 500 ms, followed by a 2.5-second blank screen. Each trial was followed by a central fixation cross with variable duration between 1 and 3 s (mean = 1.5 s). Participants rated each noun according to how often they encountered the corresponding entity or event in their daily lives, on a scale from 1 (“rarely or never”) to 3 (“often”). This familiarity judgment task was designed to encourage semantic processing of the word stimuli without emphasizing any particular semantic features or dimensions. Participants indicated their response by pressing one of three buttons on a response pad with their right hand. Stimulus presentation and response recording were performed with Psychopy 3 software (Peirce, 2007) running on a Windows desktop computer and a Celeritas fiber optic response system (Psychology Software Tools, Inc.). Stimuli were displayed on an MRI-compatible LCD screen positioned behind the scanner bore and viewed through a mirror attached to the head coil.

### MRI Data Acquisition and Processing

Images were acquired with a 3T GE Premier scanner at the Medical College of Wisconsin. Structural imaging included a T1-weighted MPRAGE volume (FOV = 256 mm, 222 axial slices, voxel size = 0.8 × 0.8 × 0.8 mm^3^) and a T2-weighted CUBE acquisition (FOV = 256 mm, 222 sagittal slices, voxel size = 0.8 × 0.8 × 0.8 mm^3^). T2*-weighted gradient-echo echoplanar images were obtained for functional imaging using a simultaneous multi-slice sequence (SMS factor = 4, TR = 1500 ms, TE = 33 ms, flip angle = 50°, FOV = 208 mm, 72 axial slices, in-plane matrix = 104 × 104, voxel size = 2 × 2 × 2 mm^3^). A pair of T2-weighted spin echo echo-planar scans (5 volumes each) with opposing phase-encoding directions was acquired before run 1, between runs 4 and 5, and after run 8, to provide estimates of EPI geometric distortion in the phase-encoding direction.

Data preprocessing was performed using fMRIPrep 20.1.0 (Esteban et al., 2018). After slice timing correction, functional images were corrected for geometric distortion, which implemented non-linear transformations estimated from the paired T2-weighted spin echo images. All images were then aligned to correct for head motion before aligning to the T1-weighted anatomical image. All voxels were normalized to have a mean of 100 and a range of 0 to 200. To optimize alignment between participants and to constrain the searchlight analysis to cortical grey matter, individual brain surface models were constructed from T1-weighted and T2-weighted anatomical data using Freesurfer and the HCP pipeline (Glasser et al., 2013). We visually checked the quality of reconstructed surfaces before carrying out the analysis. Segmentation errors were corrected manually, and the corrected images were fed back to the pipeline to produce the final surfaces. The cortex ribbon was reconstructed in standard grayordinate space with 2-mm spaced vertices, and the EPI images were projected onto this space. A general linear model was built to fit the time series of the functional data via multivariable regression. Each word (with its six presentations) was treated as a single regressor of interest and convolved with a hemodynamic response function, resulting in 320 (Experiment 1) or 300 (Experiment 2) beta coefficient maps. Regressors of no interest included head motion parameters (12 regressors), response time (z-scores), mean white matter signal, and mean cerebrospinal fluid signal. A *t* statistical map was generated for each word, and these maps were subsequently used in the searchlight RSA.

### Surface-Based Searchlight Representational Similarity Analysis

RSA was carried out using custom Python and Matlab scripts. Searchlight RSA typically employs spherical volumes moved systematically through the brain or the cortical grey matter voxels. This method, however, does not exclude signals from white matter voxels that happen to fall within the sphere, and which may contribute noise. Spherical volumes may also erroneously combine non-contiguous cortical regions across sulci. Surface-based searchlight analysis overcomes these shortcomings using circular 2-dimensional “patches” confined to contiguous vertices on the cortical surface. At each vertex, a 5-mm radius patch around the seed vertex on the midthickness surface was created, resulting in a group of vertices comprising each patch.

Representational dissimilarity matrices (RDMs) were calculated for the multimodal experiential model (the model RDM) and for each searchlight ROI (the neural RDM). Each entry in the neural RDM represented the correlation distance between fMRI responses evoked by two different words. Neural RDMs were computed for each of the 64,984 searchlight ROIs. For the model RDM, we calculated the Pearson correlation distances between each pair of words in the 65-dimensional experiential feature space. Ten additional RDMs were computed using pair-wise differences on non-semantic lexical variables, namely, number of letters, number of phonemes, number of syllables, mean bigram frequency, mean trigram frequency, orthographic neighborhood density, phonological neighborhood density, phonotactic probability for single phonemes, phonotactic probability for phoneme pairs, and word frequency (https://www.sc.edu/study/colleges_schools/artsandsciences/psychology/research_clinical_facilities/scope/). These RDMs were regressed out of the model-based RDM prior to computing the RSA correlations to remove any effects of orthographic and phonological similarity. An RDM computed from the Jaccard distance between bitmap images of the word stimuli was also used to control for low-level visual similarity between words. Spearman correlations were computed between the residual model-based RDM (after regressing out the RDMs of no interest) and the neural RDM for each ROI, resulting in a map of correlation scores on the cortical surface for each participant.

Finally, second-level analysis was performed on the correlation score maps after alignment of each individual map to a common surface template (the 32k_FS_LR mesh produced by the HCP pipeline), Fisher z-transformation, and smoothing of the maps with a 6-mm FWHM Gaussian kernel. A one-tailed, one-sample t-test against zero was applied at all vertices. FSL’s Permutation Analysis of Linear Models (PALM) was used for non-parametric permutation testing to determine cluster-level statistical inference (10,000 permutations). We used a cluster-forming threshold of z > 3.1 (p < 0.001) and a cluster-level significance level of α < 0.01. The final data were rendered on the group averaged HCP template surface.

### Partial Correlation Analyses Controlling for Unimodal Models

To test whether the cortical areas identified by the multimodal experiential model indeed encoded information about multiple sensory-motor modalities, we conducted partial correlation RSAs in which RDMs encoding the effect of a single experiential modality were partialed out, one at a time, from the full-model RDM and from the neural RDM prior to computing the correlation between the two. This analysis tested whether the multimodal model predicted the neural similarity structure of lexical concepts, at each searchlight ROI, above and beyond what could be predicted by any unimodal model. Subsets of experiential features corresponding to specific modalities were selected to form the following 7 unimodal models: (a) visual (“Color”, “Bright”, “Dark”, and “Pattern”); (b) auditory (“Sound”, “Loud”, “Low”, and “High”); (c) tactile (“Touch”, “Temperature”, ‘Weight”, and “Texture”); (d) olfactory (“Smell”); (e) gustatory (“Taste”); (f) motor (“Manipulation”, “Upper Limb”, “Lower Limb”, and “Head/Mouth”); and (g) affective (“Happy”, “Sad”, “Fearful”, and “Angry”). In these analyses, all model-based RDMs were calculated using the Euclidean distance. The RDM representing the unique contribution of a given modality (say, visual) was obtained by partialling out the RDMs based on each of the other unimodal feature subsets (e.g., auditory, tactile, olfactory, gustatory, motor, and affective) from the RDM based on that modality to create an RDM that captured modality-specific content. The residual unimodal RDM was then partialed out of the full-model and the neural RDMs used in the RSA. We conducted 7 searchlight RSAs, each controlling for the effect of a single modalilty (as well as for the lexical and visual RDMs of no interest), resulting in 7 partial correlation maps. These analyses were constrained to a region-of-interest defined by the areas in which the RSA searchlight with the full experiential model reached significance in both experiments (the red areas in Figure 2). Each partial correlation map was thresholded at an FDR-corrected P < 0.01. The conjunction of the 7 thresholded maps revealed the areas in which the multimodal model explained significant variance that was not explained by any of the unimodal models.

### Partial Correlation Analyses Controlling for Non-Experiential Models

Since previous studies have found significant RSA correlations with concept similarities computed from distributional semantics, we conducted a whole-brain RSA searchlight analysis using pairwise similarity values computed from word2vec word embeddings to verify whether it would identify the same regions found with the experiential model. We also conducted a partial correlation RSA searchlight analysis to identify areas in which the multimodal experiential model predicted the neural similarity structure of concept-related activation patterns while controlling for the similarity structure predicted by word2vec. In this analysis, significant RSA correlations indicate that multimodal experiential information accounts for a degree of similarity among neural activation patterns that is not explained by the distributional model. Word2vec similarities were computed as the cosine between the 300-dimensional word vectors trained on the Google News dataset (approximately 100 billion words), based on the continuous skip-gram algorithm and distributed by Google (https://code.google.com/archive/p/word2vec).

Finally, we evaluated the performance of the multimodal experiential model relative to the taxonomic structure of the word set (i.e., concept similarity based on membership in superordinate semantic categories) by computing mean semipartial RSA correlations averaged across all searchlight ROIs that reached significance in the main analysis. That is, searchlight RSAs were conducted for the experiential RDM after regressing out the categorical RDM and vice-versa, and the resulting RSA scores were averaged across searchlight ROIs. The difference in mean RSA scores between models was tested via permutation test with 10,000 permutations. This procedure was also used to evaluate the experiential model relative to word2vec.

### Peak Identification in the Combined Dataset Across the Two Experiments

We combined the data from the two experiments by computing the mean RSA correlation with the multimodal experiential model across all 64 participants. This allowed us to identify distinct regions with high multimodal information content across all semantic categories investigated.

## Results

### Experiment 1

The mean response rate on the familiarity judgment task was 98.7% (SD 1.9%). Intra-individual consistency in familiarity ratings across the 6 repetitions of each word was evaluated using intraclass correlations (ICCs) based on a single measurement, two-way mixed effects model and the absolute agreement definition. Results suggested generally good overall intra-individual agreement, with individual ICCs ranging from fair to excellent (mean ICC = 0.671, range: 0.438 – 0.858, all *p*s < 0.00001) (Cicchetti, 1994). To examine consistency in familiarity ratings across participants, responses to the 6 repetitions were first averaged within individuals, and the ICC across participants was calculated using the consistency definition. The resulting ICC of 0.586 (95% confidence interval [0.548, 0.625], *p* < .00001) suggested fair to good inter-individual consistency.

Group-level searchlight RSA showed a bilateral, distributed network of regions where neural similarity correlated with the semantic similarity index computed from the experiential model (Figure 1, left). In the temporal lobe, these regions included the temporal pole, superior temporal gyrus and sulcus (STG and STS), posterior middle temporal gyrus (pMTG), posterior inferior temporal gyrus (pITG), and parahippocampal gyrus (PHG), all bilaterally. The left fusiform gyrus (FG) was also involved. Parietal lobe involvement included angular and supramarginal gyri (AG and SMG), posterior superior parietal lobule (pSPL), intraparietal sulcus (IPS), precuneus (PreCun), and posterior cingulate gyrus (pCing), all bilaterally. Frontal lobe regions included the inferior, middle, and superior frontal gyri (IFG, MFG, and SFG), ventral precentral sulcus (vPreCS), rostral anterior cingulate gyrus (rACG), and orbital frontal cortex (OFC), all bilaterally. The left anterior insula was also implicated.

**Figure 1.**
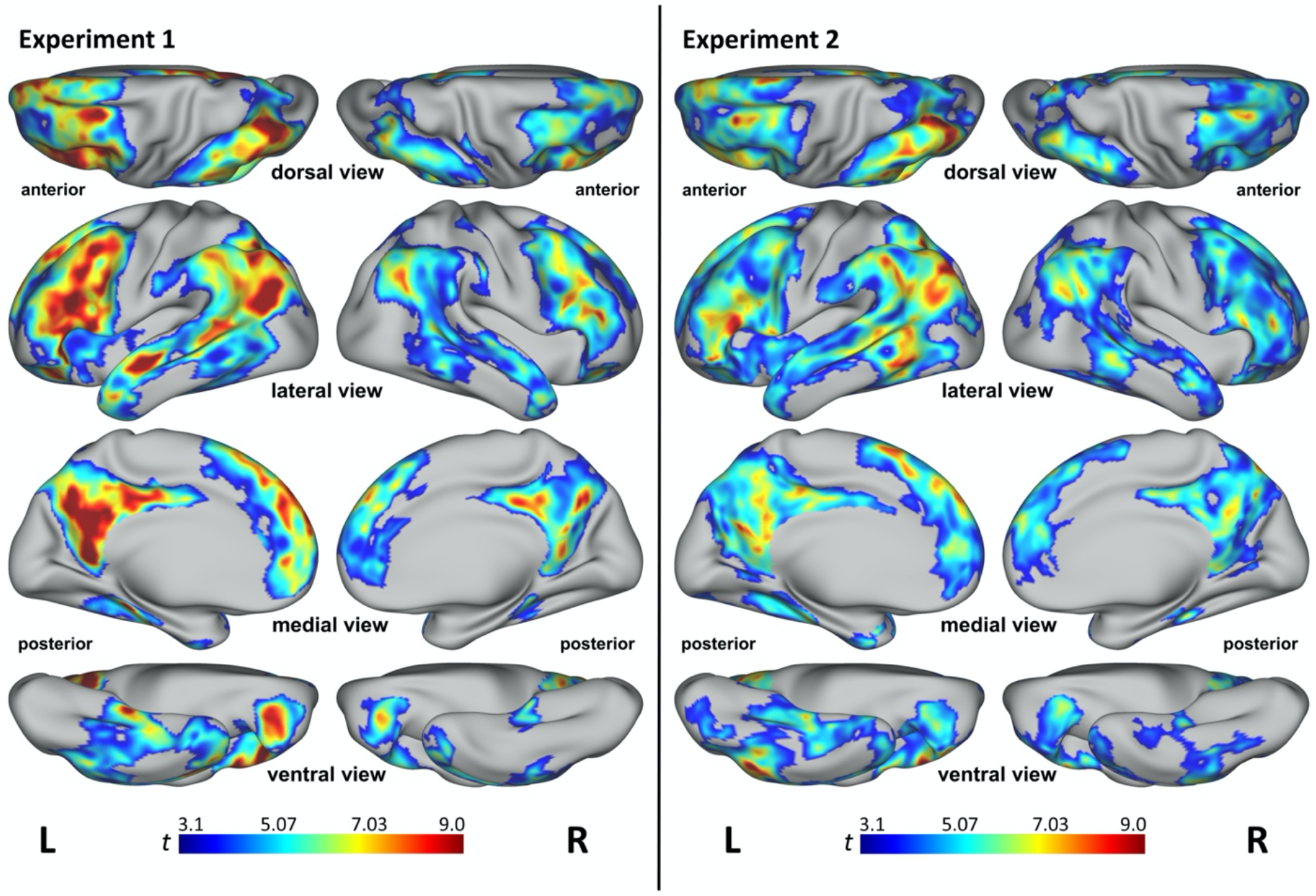
Searchlight RSA results for the multimodal experiential model from Experiment 1 (left) and Experiment 2 (right). All results are significant at p < 0.001 and cluster corrected at α < 0.01. Colors represent t values.

### Experiment 2

The mean response rate on the familiarity judgment task was 99.2% (SD = 0.63%). Intra-individual consistency analysis showed generally good overall intra-individual agreement, with individual ICCs ranging from fair to excellent (mean ICC = 0.667, range: 0.464 – 0.843, all *p*s < 0.00001) (Cicchetti, 1994). The ICC across participants was 0.533 (95% confidence interval [0.548, 0.625], *p* < .00001), suggesting fair to good inter-individual consistency.

As with Experiment 1, group-level searchlight RSA showed a bilateral, distributed network of regions where neural similarity correlated with semantic similarity as defined by the experiential model (Figure 1, right). These areas largely coincided with those identified in Experiment 1, including temporal pole, STG, STS, pMTG, pITG, FG, PHG, AG, SMG, IPS, pSPL, pCing, PreCun, IFG, MFG, SFG, vPreCS, rACG, and OFC, all bilaterally; and left anterior insula. Areas of overlap between the two experiments are shown in red in Figure 2. The percentage of overlapping vertices between the two experiments, as measured by the Jaccard index, was 73.2%.

**Figure 2.**
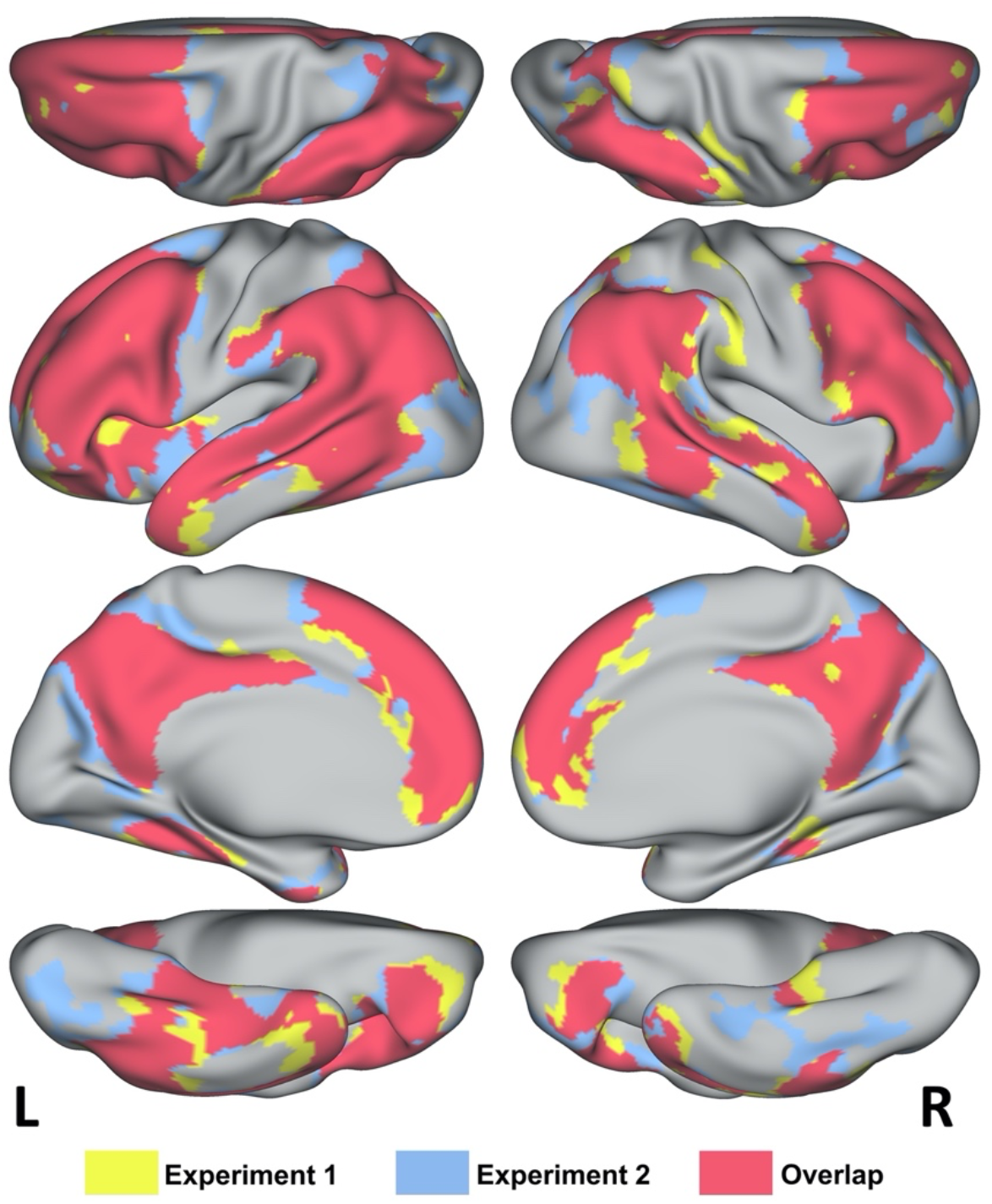
Cortical areas where the RSA score for the multimodal experiential model reached significance in Experiment 1 (yellow), in Experiment 2 (blue), or in both experiments (red). Figure 2-1 shows that the areas in the overlap region also reached significance when RSA for the multimodal model was controlled for the effects of unimodal experiential models.

### Partial correlation analyses controlling for unimodal models

In both experiments, we found that all cortical areas detected in the main analysis showed significant RSA correlations for the RDM based on the multimodal model after controlling for the effects of each unimodal experiential model (Figure 2-1). These results indicate that the relationships observed between the multimodal experiential model and neural similarity patterns in these heteromodal regions could not be fully explained by any of the unimodal models.

**Figure 2-1.**
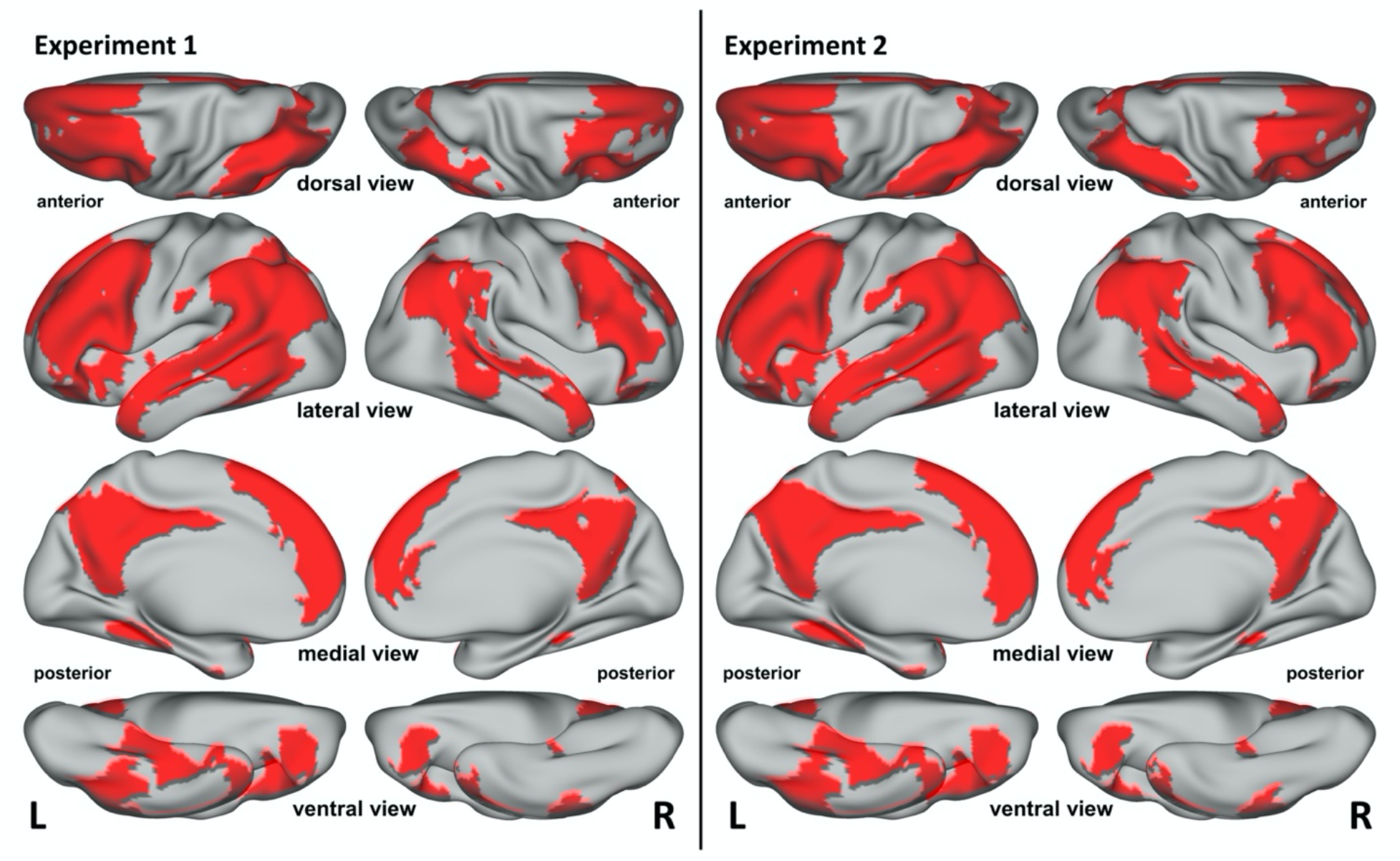
RSA results for the multimodal model after removing contributions from each of 7 unimodal models representing visual, auditory, somatosensory, smell, taste, action, and affective experiential content. Shown in red are the regions within the overlap map from Figure 2 where the multimodal model explained additional variance compared to all of the unimodal models in isolation.

### Partial correlation analyses controlling for non-experiential models

Visual inspection of the representational similarity matrices for the multimodal experiential and word2vec models (Figure 3-1) shows that both models reflect the taxonomic structure of lexical concepts to some extent. In Experiment 1, searchlight RSA for the word2vec model revealed a set of cortical areas similar to those found for the multimodal experiential model, although with substantially less extensive clusters (Figure 3-2). In Experiment 2, the results for word2vec were very similar to those for the experiential model. In both experiments, we found that, even when word2vec similarity was controlled for, the multimodal experiential model predicted the neural similarity pattern for lexical concepts in the same regions found in the main analysis (Figure 3). The extent of the clusters was appreciably reduced relative to the main analysis only in Experiment 2, particularly in the right hemisphere.

**Figure 3.**
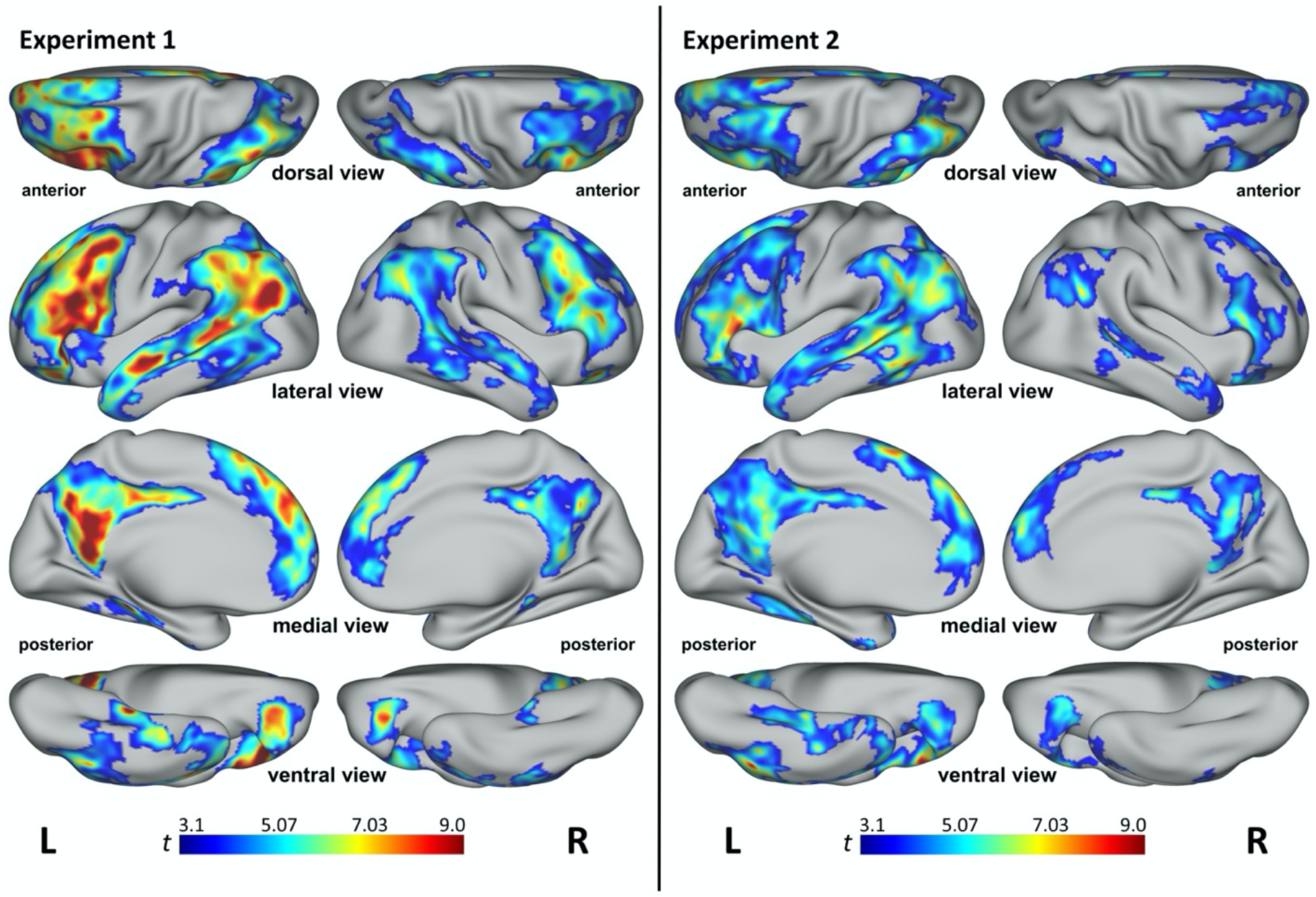
Searchlight RSA results for the multimodal experiential model after controlling for word2vec similarity from Experiments 1 (left) and 2 (right). This analysis identified areas in which the experiential model accounted for patterns of neural similarity that were not explained by word2vec. All results are significant at p < 0.001 and cluster corrected at α < 0.01. Colors represent t values. Figure 3-1 (Extended Data) shows the representational similarity matrices for the multimodal experiential and word2vec models with words sorted according to superordinate categories. Figure 3-2 shows the searchlight RSA results for the word2vec model. Figure 3-3 shows the unique prediction performance of the multimodal experiential model relative to the word2vec model and vice-versa, averaged across searchlight patches. Figure 3-4 shows the unique prediction performance of the multimodal experiential model relative to the categorical model and vice-versa, averaged across searchlight patches.

**Figure 3-1:**
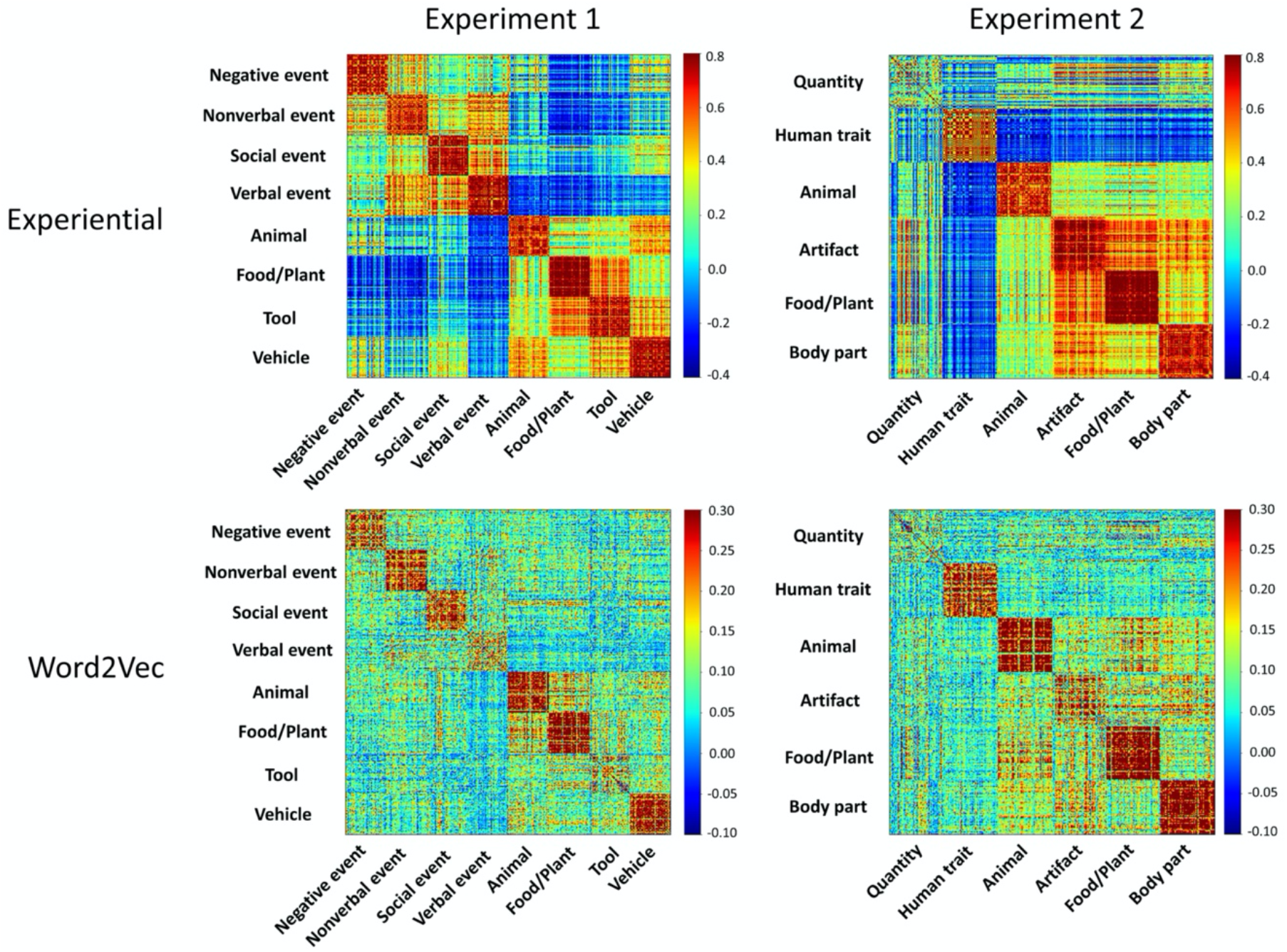
Representational similarity matrices for the multimodal experiential and word2vec models in Experiment 1 (left) and Experiment 2 (right), with words sorted by superordinate categories. Both models capture categorical structure to varying degrees depending on category, as demonstrated by higher similarity values (red) for item pairs within categories. The “Quantity” items in Experiment 2, which included concepts of time, distance, size, area, volume, amplitude, etc., do not appear to form a coherent category in either model.

**Figure 3-2:**
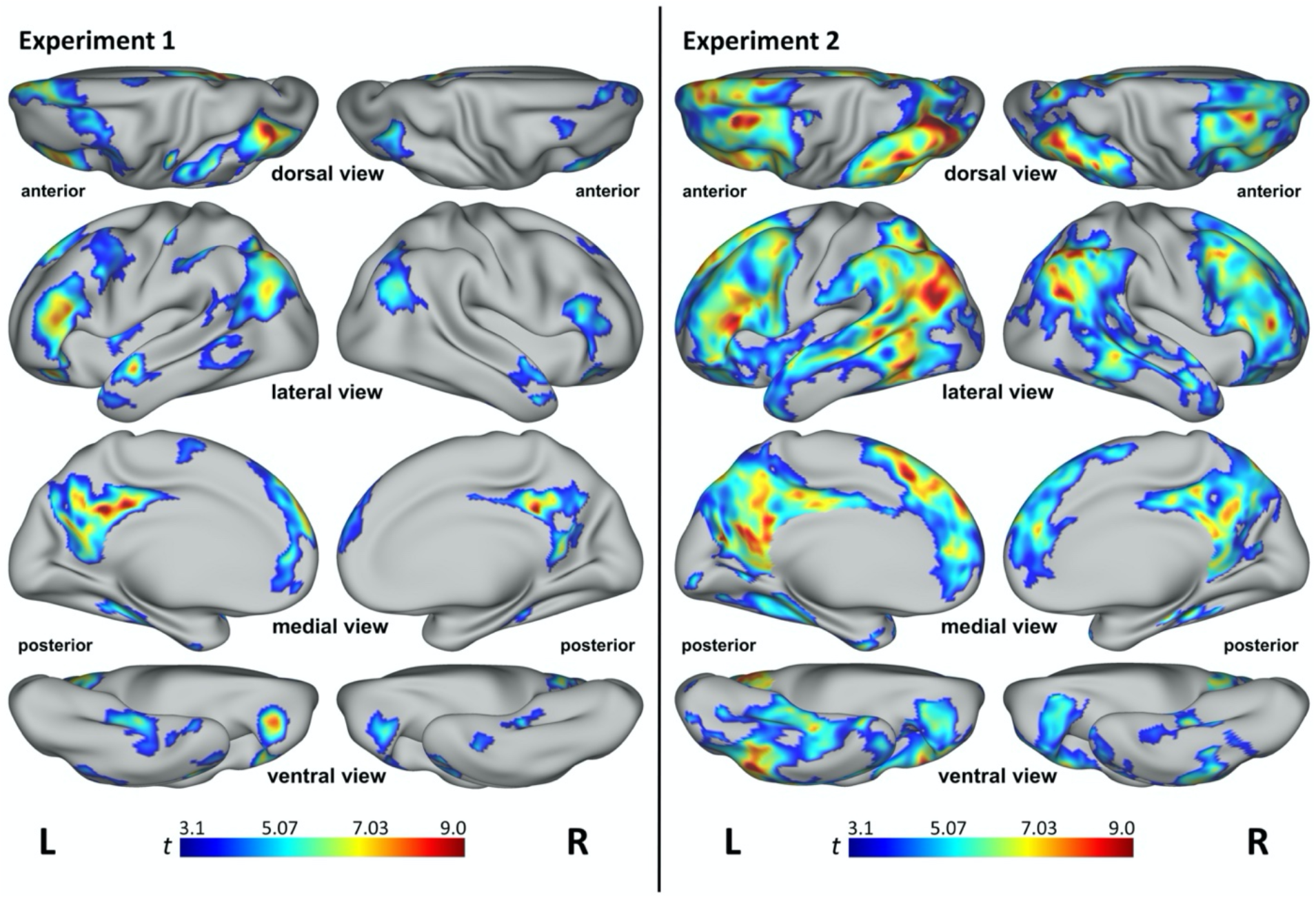
Searchlight RSA results for the word2vec model from Experiments 1 (left) and 2 (right).

The mean semipartial RSAs comparing the performances of the two models showed that, in both experiments, the neural similarity structure of concepts was significantly closer to the similarity structure predicted by the multimodal experiential model than that predicted by word2vec (p < 0.0001; Figure 3-3). Similarly, in both experiments, mean semipartial RSAs showed a significant advantage for the experiential model compared to the categorical structure model (p < 0.0001; Figure 3-4).

**Figure 3-3:**
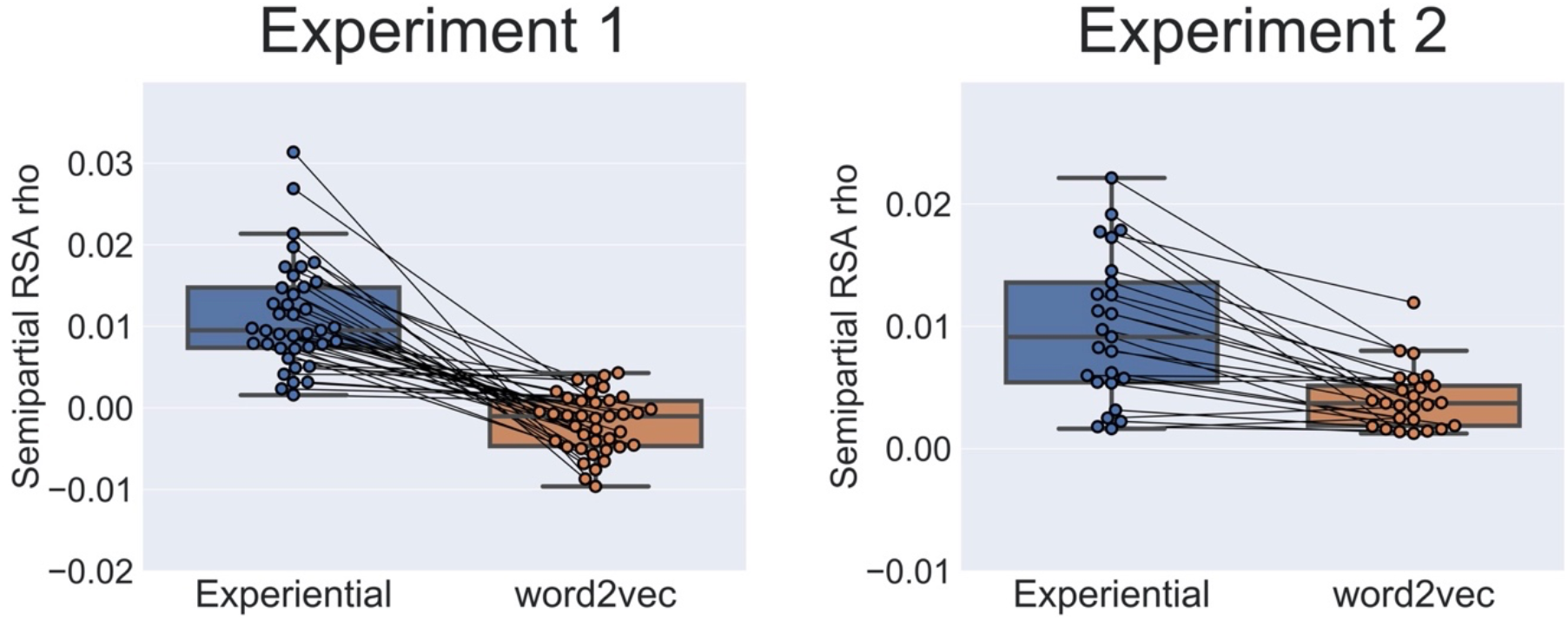
Unique prediction performance of the multimodal experiential model and of the word2vec model relative to each other. The boxplots show the mean semipartial RSA correlation for the experiential model controlling for the word2vec model (blue) and vice-versa (orange), averaged across all searchlight ROIs that reached significance in the main analysis (areas depicted in red in Figure 2). Each paired data point corresponds to one participant. Both experiments showed a significant advantage for the experiential model (p < 0.0001, permutation test).

**Figure 3-4:**
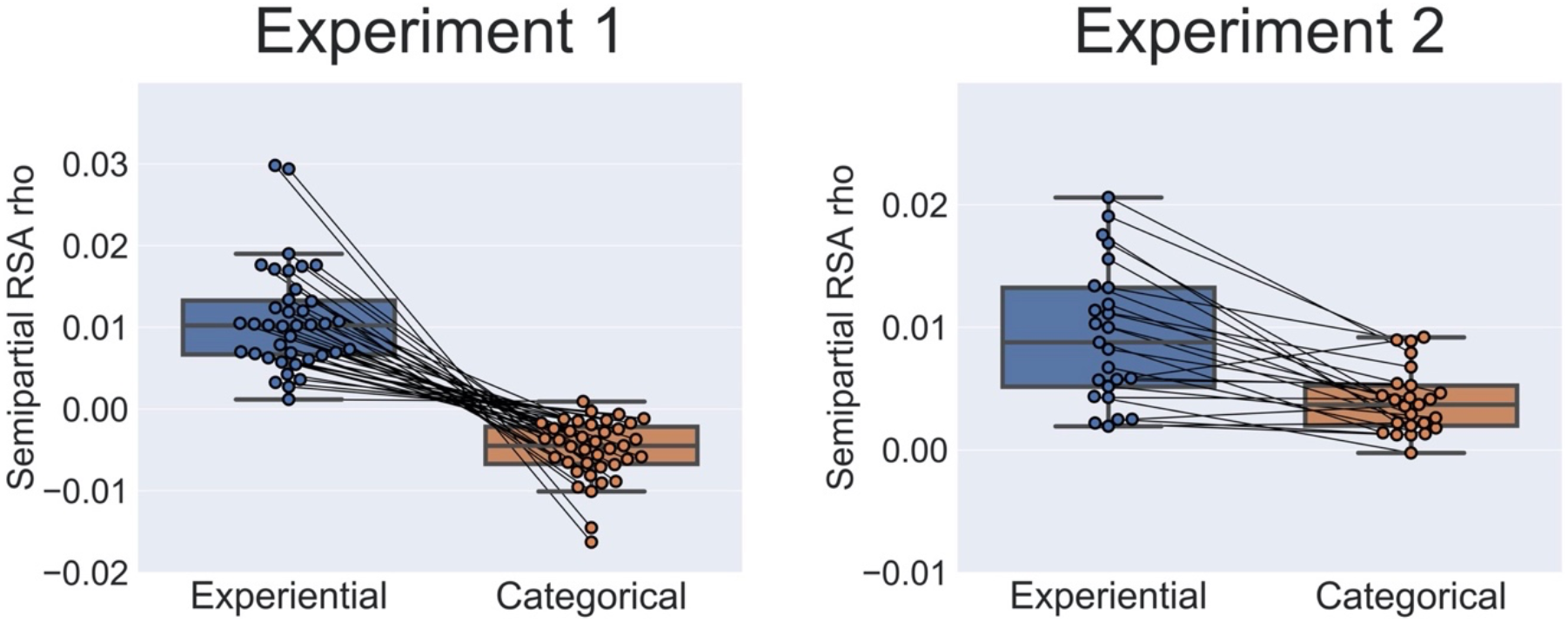
Unique prediction performance of the multimodal experiential model and of the categorical model relative to each other. The boxplots show the mean semipartial RSA correlation for the experiential model controlling for the categorical model (blue) and vice-versa (orange), averaged across all searchlight ROIs that reached significance in the main analysis (areas depicted in red in Figure 2). Each paired data point corresponds to one participant. Both experiments showed a significant advantage for the experiential model (p < 0.0001, permutation test).

### Peak identification in the combined dataset

The combined RSA searchlight map for the experiential model, averaged across participants from both experiments, clearly shows the existence of distinct regions of high multimodal information content within the semantic network (Figure 4). In the left hemisphere, local peaks were evident in the IFG, posterior MFG, posterior superior frontal sulcus (SFS), mid-SFG, ventral PreCS, lateral OFC, AG, pIPS, pMTG/pSTS, PreCun, pCing, retrosplenial cortex (RSC), anterior STS, lateral temporal pole, PHG and anterior FG, with smaller peaks in similar right hemisphere regions (Table 1).

**Figure 4.**
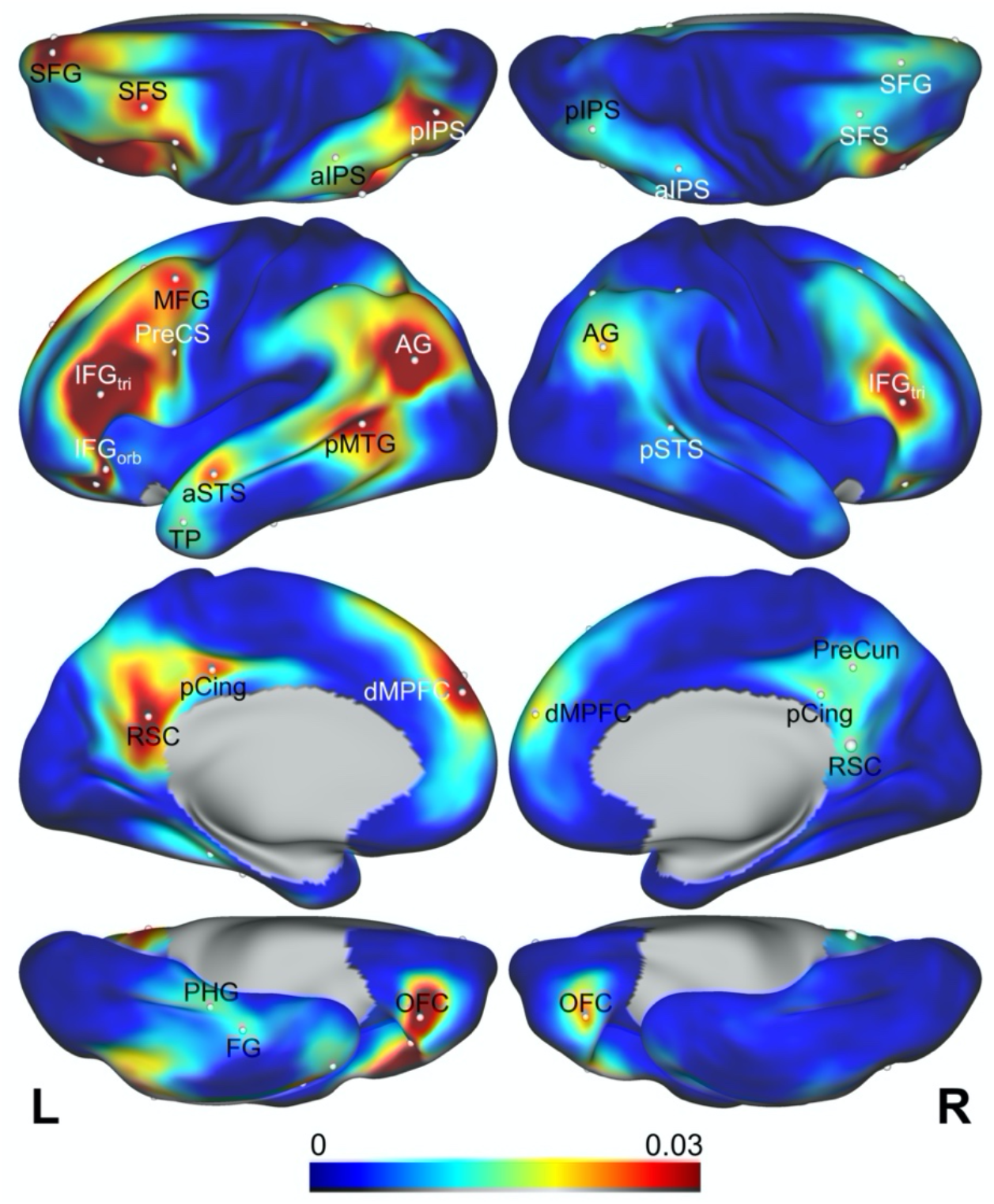
RSA scores for the multimodal experiential model averaged across all participants in Experiments 1 and 2 (n = 64). The RSA peaks reported in Table 1 are indicated.

**Table 1.**
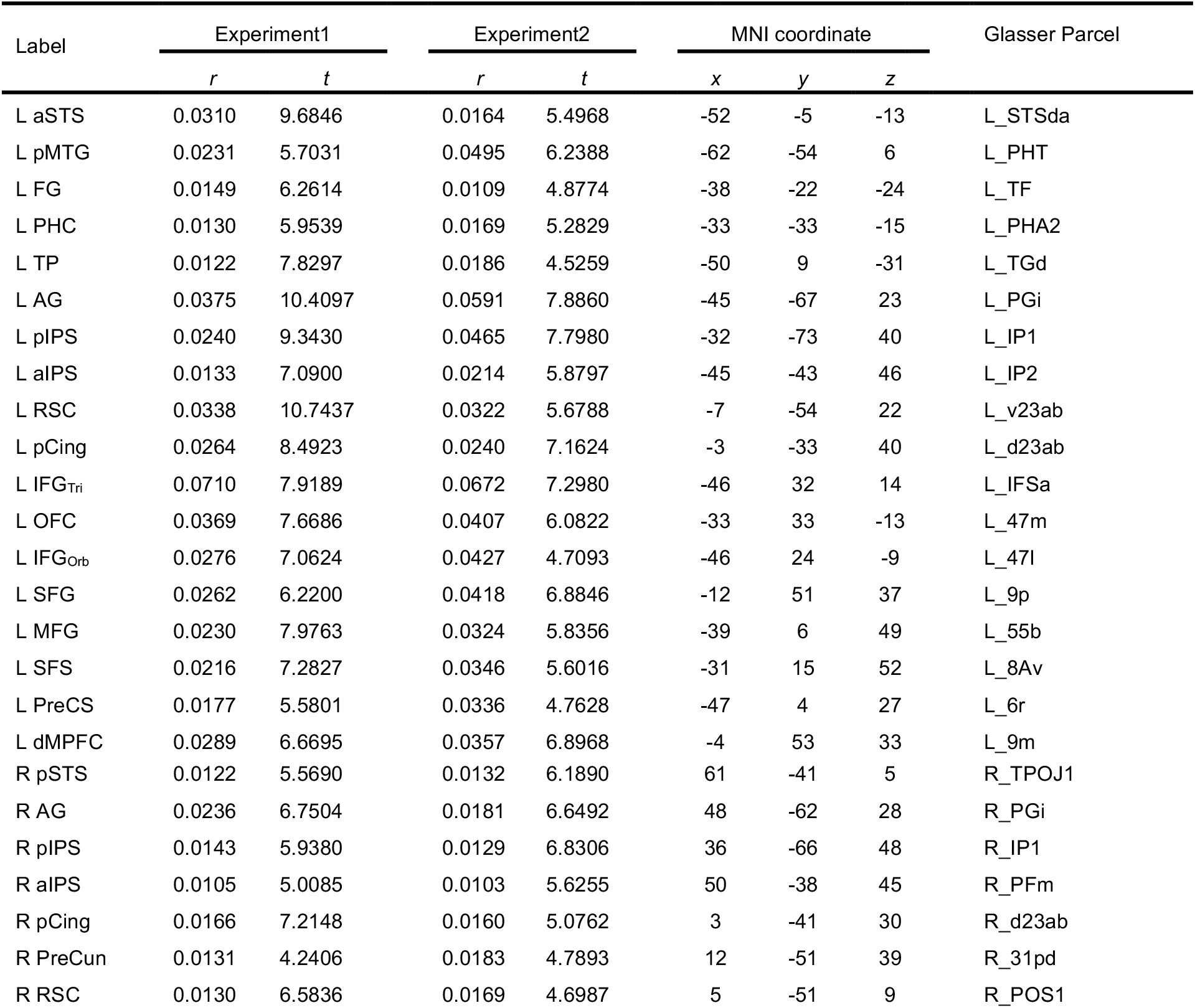
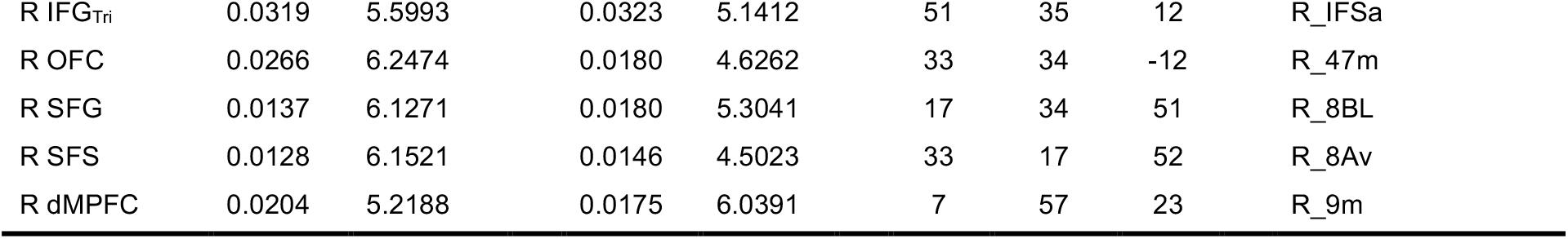
Peaks in the combined RSA correlation map, averaged across all participants from both experiments. Peak labels correspond to those used in Figure 4. Coordinates refer to the MNI 152 2009 template space. Parcel labels are taken from the surface-based atlas in Glasser et al. (2016). AG: angular gyrus; aIPS: anterior intraparietal sulcus; aSTS: anterior superior temporal sulcus; FG: fusiform gyrus; IFGOrb: pars orbitalis of the inferior frontal gyrus; IFGTri: pars triangularis of the inferior frontal gyrus; MFG: middle frontal gyrus; OFC: orbitofrontal cortex; pCing: posterior cingulate cortex; PHC: parahippocampal cortex; pIPS: posterior intraparietal sulcus; pMTG: posterior middle temporal gyrus; preCS: precentral sulcus; preCun: precuneus; RSC: retrosplenial cortex; SFG: superior frontal gyrus; SFS: superior frontal sulcus; TP: temporal pole.

| Label | Experiment1 |  | Experiment2 |  | MNI coordinate |  |  | Glasser Parcel |
| --- | --- | --- | --- | --- | --- | --- | --- | --- |
|  | *r* | *t* | *r* | *t* | *x* | *y* | *z* |  |
| L aSTS | 0.0310 | 9.6846 | 0.0164 | 5.4968 | -52 | -5 | -13 | L_STSda |
| L pMTG | 0.0231 | 5.7031 | 0.0495 | 6.2388 | -62 | -54 | 6 | L_PHT |
| L FG | 0.0149 | 6.2614 | 0.0109 | 4.8774 | -38 | -22 | -24 | L_TF |
| L PHC | 0.0130 | 5.9539 | 0.0169 | 5.2829 | -33 | -33 | -15 | L_PHA2 |
| L TP | 0.0122 | 7.8297 | 0.0186 | 4.5259 | -50 | 9 | -31 | L_TGd |
| L AG | 0.0375 | 10.4097 | 0.0591 | 7.8860 | -45 | -67 | 23 | L_PGi |
| L pIPS | 0.0240 | 9.3430 | 0.0465 | 7.7980 | -32 | -73 | 40 | L_IP1 |
| L aIPS | 0.0133 | 7.0900 | 0.0214 | 5.8797 | -45 | -43 | 46 | L_IP2 |
| L RSC | 0.0338 | 10.7437 | 0.0322 | 5.6788 | -7 | -54 | 22 | L_v23ab |
| L pCing | 0.0264 | 8.4923 | 0.0240 | 7.1624 | -3 | -33 | 40 | L_d23ab |
| L IFG<sub>Tri</sub> | 0.0710 | 7.9189 | 0.0672 | 7.2980 | -46 | 32 | 14 | L_IFSa |
| L OFC | 0.0369 | 7.6686 | 0.0407 | 6.0822 | -33 | 33 | -13 | L_47m |
| L IFG<sub>Orb</sub> | 0.0276 | 7.0624 | 0.0427 | 4.7093 | -46 | 24 | -9 | L_47l |
| L SFG | 0.0262 | 6.2200 | 0.0418 | 6.8846 | -12 | 51 | 37 | L_9p |
| L MFG | 0.0230 | 7.9763 | 0.0324 | 5.8356 | -39 | 6 | 49 | L_55b |
| L SFS | 0.0216 | 7.2827 | 0.0346 | 5.6016 | -31 | 15 | 52 | L_8Av |
| L PreCS | 0.0177 | 5.5801 | 0.0336 | 4.7628 | -47 | 4 | 27 | L_6r |
| L dMPFC | 0.0289 | 6.6695 | 0.0357 | 6.8968 | -4 | 53 | 33 | L_9m |
| R pSTS | 0.0122 | 5.5690 | 0.0132 | 6.1890 | 61 | -41 | 5 | R_TPOJ1 |
| R AG | 0.0236 | 6.7504 | 0.0181 | 6.6492 | 48 | -62 | 28 | R_PGi |
| R pIPS | 0.0143 | 5.9380 | 0.0129 | 6.8306 | 36 | -66 | 48 | R_IP1 |
| R aIPS | 0.0105 | 5.0085 | 0.0103 | 5.6255 | 50 | -38 | 45 | R_PFm |
| R pCing | 0.0166 | 7.2148 | 0.0160 | 5.0762 | 3 | -41 | 30 | R_d23ab |
| R PreCun | 0.0131 | 4.2406 | 0.0183 | 4.7893 | 12 | -51 | 39 | R_31pd |
| R RSC | 0.0130 | 6.5836 | 0.0169 | 4.6987 | 5 | -51 | 9 | R_POS1 |
| R IFG<sub>Tri</sub> | 0.0319 | 5.5993 | 0.0323 | 5.1412 | 51 | 35 | 12 | R_IFSa |
| R OFC | 0.0266 | 6.2474 | 0.0180 | 4.6262 | 33 | 34 | -12 | R_47m |
| R SFG | 0.0137 | 6.1271 | 0.0180 | 5.3041 | 17 | 34 | 51 | R_8BL |
| R SFS | 0.0128 | 6.1521 | 0.0146 | 4.5023 | 33 | 17 | 52 | R_8Av |
| R dMPFC | 0.0204 | 5.2188 | 0.0175 | 6.0391 | 7 | 57 | 23 | R_9m |

## Discussion

We sought to clarify the large-scale architecture of the concept representation system by identifying cortical regions whose activation patterns encode multimodal experiential information about individual lexical concepts. Across two independent experiments, each involving a large number and a wide range of concepts (for a total of 522 unique lexical concepts), we detected multimodal concept representation in widespread heteromodal cortical regions, bilaterally, including anterior, posterior, and ventral temporal cortex, inferior and superior parietal lobules, medial parietal cortex, and medial, dorsal, ventrolateral, and orbital frontal regions. In all of these areas, the multimodal experiential model accounted for significant variance in the neural similarity structure of lexical concepts that could not be explained by semantic structure derived from unimodal experiential models, categorical models, or word co-occurrence patterns. These results confirm and extend previous neuroimaging studies indicating that the concept representation system is highly distributed and that multimodal experiential information is encoded throughout the system. The present study identified several distinct regions displaying relatively high conceptual information content, including regions not typically associated with lexical semantic processing, such as the orbitofrontal cortex, superior frontal gyrus, precuneus, and posterior cingulate gyrus. We propose that these regions with strong multimodal information content are good candidates for the convergence zones postulated by certain models of concept representation and retrieval (Damasio, 1989; Mesulam, 1998; Meyer and Damasio, 2009).

The network of brain regions identified in the current study closely resembles the network identified previously in a meta-analysis of 120 functional imaging studies on semantic processing (Binder et al., 2009). It has been argued that some of these regions, such as the angular gyrus, do not actually contribute to conceptual processing, only appearing to be activated in semantic tasks due to differences in task difficulty (e.g., Humphreys et al., 2021). The present results provide direct evidence that all brain regions highlighted in the previous meta-analysis indeed represent conceptual information during semantic word processing. Unlike previous studies based on cognitive subtraction paradigms, our RSA results cannot be explained by systematic differences in difficulty or task requirements.

In contrast to previous RSA studies of concept representation (Anderson et al., 2015; Carota et al., 2021; Devereux et al., 2013; Martin et al., 2018), the network identified in the present study includes extensive cortex in the anterior temporal lobe, a region strongly implicated in high-level semantic representation (Lambon Ralph et al., 2017). While the present study did not show significant RSA correlations in the anterior ventrolateral temporal lobes, it is important to note that these regions (as well as the ventromedial prefrontal cortex), exhibit typically low BOLD signal-to-noise ratios owing to magnetic susceptibility effects. Therefore, the present data do not allow us to derive conclusions about conceptual information content in these areas.

Further studies using echo-planar imaging parameters optimized for detecting BOLD signal in these areas are needed to address this issue.

The concept representation network identified in the current study also closely resembles the set of cortical regions referred to as the “default mode network” (DMN) (Buckner et al., 2008). Several functional connectivity studies indicate that these areas function as hubs, or convergence zones, for the multimodal integration of sensory-motor information (Buckner et al., 2009; Sepulcre et al., 2012; Heuvel and Sporns, 2013; Margulies et al., 2016; Murphy et al., 2018). They have been shown to be equidistant from primary sensory and motor areas in “stepwise” functional connectivity analyses, sitting at the end of the principal gradient of cortical connectivity going from unimodal to heteromodal areas. Activity in these areas has also been shown to correlate positively with the relevance of multiple sensory-motor features of word meaning (Fernandino et al., 2016a) and to be associated with the level of experiential detail present in ongoing thought (Smallwood et al., 2016; Sormaz et al., 2018). The current results provide novel evidence that activity patterns in DMN regions also reflect conceptual content conveyed by individual words. This further supports the view that concept retrieval is a major component of the brain’s “default mode” of processing (Binder et al., 1999, 2009; Andrews-Hanna et al., 2014; Yeshurun et al., 2021).

Our results confirm and extend previous RSA studies that identified portions of this network using semantic models and word stimuli. Three studies implicated anteromedial temporal cortex, particularly perirhinal cortex, as a semantic hub (Bruffaerts et al., 2013; Liuzzi et al., 2015; Martin et al., 2018). All used semantic models based on crowd-sourced feature production lists, and all used a feature verification task during fMRI (e.g., “WASP – Does it have paws?”). Prior studies combining searchlight RSA with either taxonomic (Devereux et al., 2013; Carota et al., 2021) or distributional (Anderson et al., 2015; Carota et al., 2021) semantic models have implicated more widespread regions, including posterior lateral temporal cortex, inferior parietal lobe, posterior cingulate gyrus, and prefrontal cortex. The two studies using taxonomic models (Devereux et al., 2013; Carota et al., 2021) showed similar involvement of the left posterior STS and MTG, with extension into adjacent AG and SMG. In contrast, the two studies using distributional models (Anderson et al., 2015; Carota et al., 2021) found little or no posterior temporal involvement, and inferior parietal involvement was confined mainly to the left SMG. Frontal cortex involvement was uniformly present but highly variable in extent and location across those studies. Two studies reported involvement of the posterior cingulate/precuneus (Anderson et al., 2015; Devereux et al., 2013).

Several factors may have negatively impacted sensitivity and reliability in those studies. First, ROI-based RSAs show that, relative to experiential models of concept representation, taxonomic and distributional models are consistently less sensitive to the neural similarity structure of lexical concepts (Fernandino et al., 2022). Furthermore, most of the prior studies used volume-based spherical searchlights, which typically sample a mix of grey and white matter voxels, while the surface-based approach used in the present study ensures that only contiguous cortical gray matter voxels are included, thus reducing noise from uninformative voxels. Finally, the nature of the task and the particularities of the concept set used as stimuli can affect both the sensitivity of the analysis and the cortical distribution of the RSA searchlight map, and variations in these properties may underlie some of the variation in results across studies. We dealt with this last issue by (1) employing large numbers of concepts from diverse semantic categories and (2) analyzing data from two independent experiments to identify areas displaying reliable representational correspondence with the semantic model across different concept sets and different participant samples.

The finding of extensive frontal lobe involvement in concept “representation” deserves comment. Studies of brain damaged individuals and functional imaging experiments in the healthy brain have long been interpreted as supporting the classic view that ascribes to frontal cortex an executive control rather than an information storage function in the brain (Stuss and Benson, 1986; Kimberg and Farah, 1993; Thompson-Schill et al., 1997; Wagner et al., 2001). Nevertheless, nearly all RSA studies of concept representation have observed similarity structure correlations in prefrontal regions. While these observations do not speak directly to the distinction between “storage” and “control” of information, we believe they can be reconciled with the classic view by postulating a fine-grained organization of control systems, in which prefrontal cortex is tuned, at a relatively small scale, to particular sensory-motor and affective features. Neurophysiological studies in nonhuman primates provide evidence for tuning of prefrontal neurons to preferred stimulus modalities (Romanski, 2007), as well as differential connectivity across the prefrontal cortex with various sensory systems (Barbas and Mesulam, 1981; Petrides, 2005). A few human functional imaging studies provide similar evidence for sensory modality tuning in prefrontal cortex (Greenberg et al., 2010; Michalka et al., 2015; Tobyne et al., 2017). If conceptual representation in temporal and parietal cortex is inherently organized according to experiential content, it seems plausible that controlled activation and short-term maintenance of this information would require similarly fine-grained control mechanisms. We propose that the information represented in these prefrontal regions reflects their entrainment to experiential representations stored primarily in temporoparietal cortex, providing context-dependent control over their level of activation.

Related to this issue is the question of how similar the many regions identified by RSA are to each other in terms of their representational structure. Although RSA ensures that the neural similarity structure of all these regions is related to the similarity structure encoded in the semantic model, representational structure should be expected to vary to some degree across distinct functional regions, given their unique computational properties and connectivity profiles. More research is needed to investigate potential regional differences in representational content.

## Acknowledgements

This work was supported by National Institute on Deafness and Other Communication Disorders (NIDCD) grant R01 DC016622, by the Intelligence Advanced Research Projects Activity under Grant FA8650-14-C-7357, and by a grant from the Advancing a Healthier Wisconsin Foundation (Project #5520462). The authors thank Volkan Arpinar, Elizabeth Awe, Joseph Heffernan, Steven Jankowski, Jedidiah Mathis, and Megan LeDoux for technical assistance, as well as three anonymous reviewers for helpful comments and suggestions on a previous version of this article.

